# Context dependent roles for RB-E2F transcriptional regulation in tumor suppression

**DOI:** 10.1101/399956

**Authors:** Michael J. Thwaites, Matthew J. Cecchini, Daniel T. Passos, Frederick A. Dick

## Abstract

RB-E2F transcriptional control plays a key role in regulating the timing of cell cycle progression from G1 to S-phase in response to growth factor stimulation. Despite this role, it is genetically dispensable for cell cycle exit in primary fibroblasts in response to growth arrest signals. Mice engineered to be defective for RB-E2F transcriptional control at cell cycle genes were also found to live a full lifespan with no susceptibility to cancer. Based on this background we sought to probe the vulnerabilities of RB-E2F transcriptional control defects found in *Rb1*^*R461E,K542E*^ mutant mice (*Rb1*^*G*^) through genetic crosses with other mouse strains. We generated *Rb1*^*G/G*^ mice in combination with *Trp53* and *Cdkn1a* deficiencies, as well as in combination with Kras^G12D^. The *Rb1*^*G*^ mutation enhanced *Trp53* cancer susceptibility, but had no effect in combination with *Cdkn1a* deficiency or Kras^G12D^. Collectively, this study indicates that compromised RB-E2F transcriptional control is not uniformly cancer enabling, but rather has potent oncogenic effects when combined with specific vulnerabilities.

## Introduction

The maintenance of cell cycle control is crucial to the normal development and homeostasis of multicellular organisms (1). In addition, misregulation of the cell cycle is widespread in tumorigenesis (2). To ensure that cells only replicate their genome once per cell cycle, the regulation of G1 to S-phase is tightly controlled (3). At the core of G1-S regulation are Cyclin dependent kinases (CDKs) and the Retinoblastoma (RB) family of proteins. Proliferative signals generally activate Ras and lead to Cyclin D-CDK4 or 6 upregulation, phosphorylation of RB, and the release of activator E2F transcription factors to induce cell cycle entry (4). This is complemented by CDK phosphorylation of the RB family protein p130 that disassembles the DREAM transcriptional repressor complex, further contributing to E2F activation in early G1 (5). In addition, Cyclin E-CDK2 is negatively regulated by the CDK inhibitor protein p27 in late G1 and its degradation coincides with maximal CDK2 activation and the commitment to S-phase entry (6). Thus, both CDKs and RB family members are key to the commitment step to enter the cell cycle and over expression of G1 Cyclin-CDKs accelerates entry into S-phase, as does loss of RB, or the combination of its family members p107 and p130 (7-9). While CDK and E2F regulation are well known in cell cycle control, emerging roles in cell lineage commitment suggest that RB-E2F transcription may serve more purposes than just cell cycle entry decisions, as it is only one piece of a complex E2F transcriptional network that operates in the G1 phase (10).

In addition to regulating entry into the cell cycle, many of the same molecules function to execute a transient cell cycle arrest, or more permanent cell cycle exit decisions. For example, DNA damage stabilizes p53 and leads to transcriptional activation of the CDK inhibitor p21 (11). In S-phase this inhibits CDK2 and blocks cell cycle progression, while protein phosphatases dephosphorylate and activate RB family members (12). RB is genetically required for cell cycle exit in response to DNA damage (13), while combined deficiency of p107 and p130 does not affect this cell cycle decision (13). However, kinetic experiments suggest that transcriptional repression of E2F target genes may be too slow in comparison with the inhibition of DNA synthesis to explain RB’s mechanism of arrest (14). In addition to regulating E2Fs, RB is also capable of stabilizing the CDK inhibitor p27 through the direct inactivation of Skp2 (14, 15). Thus, RB also contributes to a transcription independent mechanism of CDK regulation to arrest the cell cycle. This raises the question of how RB-E2F regulation fits into the complex network of CDK inhibition, and RB-family mediated transcriptional control, that contributes to cell cycle arrest and RB’s role as a tumor suppressor.

To determine the contexts where RB-E2F transcriptional control is most critical, we established a genetically modified mouse line in which the endogenous RB protein is engineered to possess substitutions that interfere with RB binding to the transactivation domain of E2F proteins (16, 17). These mice (called *Rb1*^*G*^) are viable, fertile, and are not cancer prone, but they possess a partially penetrant muscle wasting phenotype that compromises survival of some neonatal animals. In order to determine how the cell cycle is regulated in cells from these animals, we crossed *Rb1*^*G/G*^ mice with *Cdkn1b*^*−/−*^ to test the additive effect of losing CDK inhibition by p27 (18). Cells from *Rb1*^*G/G*^; *Cdkn1b*^*−/−*^ double mutants possess a synthetic DNA damage induced cell cycle arrest defect that neither mutant possesses alone (18). In addition, these mice are highly cancer prone and succumb to pituitary tumors as seen in *Rb1*^*+/−*^ mice. This work suggests that RB-E2F transcriptional control and CDK inhibition by p27 are at least partially redundant in cell cycle control and tumor suppression. In an effort to extend this analysis and better understand the role of RB-E2F transcriptional regulation we crossed *Rb1*^*G/G*^ mice with strains deficient for p53 and p21, as well as with a strain that expresses an activated form of Kras. The RB-E2F regulatory defect enhanced cancer susceptibility of *Trp53*^*−/−*^ mice, but had no effect in combination with *Cdkn1a* deficient animals. Lastly, activation of Kras^G12D^ using and UBC9 driven CreERT2 resulted in benign hyperplastic growths, and Kras^G12D^ in *Rb1*^*G/G*^ mice failed to result in a more severe form than activation of Kras alone. Taken together these experiments indicate that defective RB-E2F transcriptional control has potent oncogenic effects in combination with specific mutations in other genes, but is not uniformly cancer promoting.

## Materials and Methods

### Ethics Statement

All animals were housed and handled as approved by the UWO animal care committee (protocol 2016-038) and Canadian Council on Animal Care (CCAC) guidelines. All animals were euthanized when they had reached ethical endpoints based on activity levels, grooming, and body weight.

### Cell cycle analysis

Mouse embryonic fibroblasts (MEFs) were derived from E13.5 embryos of the indicated genotypes as previously described (19). Cells were cultured using standard methods in Dulbecco’s modified Eagle’s medium containing 10% fetal bovine serum (FBS), 2mM glutamine, 50U/ml penicillin, and 50μg/ml streptomycin. For serum starvation, cells were cultured in media containing 0.1% FBS for 60 hours before harvesting or restimulating with 10% serum. To induce cell cycle arrest, cells were exposed to 15 Gy of ionizing radiation as previously described (18). 48 hours after treatment cells were labeled with BrdU for 2 hours. For all experiments cell cycle analysis by propidium iodide and BrdU staining was then carried out as previously reported (20).

### Generation and analysis of experimental animals

*Trp53*^*−/−*^ mice (129-Trp53^tm^1^Tyj^/J), and *Cdkn1a*^*−/−*^ mice (B6.129S6(Cg)-Cdkn1a^tm^1^Led^/J) have been described previously and were obtained from Jackson Laboratory (21, 22). These two mutations were combined with the *Rb1*^*R461E,K542E*^ (*Rb1*^*G*^) mouse model and were genotyped as previously described (17, 21, 22). Mice were monitored for tumor development and sacrificed at animal protocol endpoints. Survival data were subjected to Kaplan-Meier analysis, and significant differences were compared using a log rank test.

*LSL-Kras*^*G12D*^ mice (B6.129S4-Kras^tm^4^Tyj^/J) mice and *UBC-Cre-ERT2* (B6.Cg-Tg(UBC-cre/ERT2)1Ejb/2J) were combined with our *Rb1*^*G/G*^ mouse model to produce both control *Kras*^*G12D*^; *Cre-ERT2* animals and experimental *Rb1*^*G/G*^; *Kras*^*G12D*^; *Cre-ERT2* animals (17, 23, 24). Animals were then injected with 75mg/kg tamoxifen at 8 weeks of age resulting in oncogenic *Kras*^*G12D*^ expression sporadically throughout their body. Animals were then monitored for tumor formation and sacrificed at endpoints. Survival data were subjected to Kaplan-Meier analysis, and significant differences were compared using a log rank test.

### Histological analysis of tumors

Following euthanasia, mice were subject to necropsy where tissues of interest were fixed in formaldehyde for 72 hours. Tissues were then washed twice in PBS before storage in 70% ethanol. Tissues were then embedded in paraffin, 5μm sections were then cut and stained with hematoxylin and eosin (H&E). Images were obtained using a Zeiss Axioskop 40 microscope and Spot Flex camera and software (Mississauga, Ontario, Canada).

## Results

### Defective RB-E2F transcriptional control results in accelerated cell cycle entry

We previously reported the generation of a mouse strain in which the endogenous *Rb1* gene was modified to express a synthetic mutant RB protein that is designed to disrupt RB’s ability to interact with the activation domain of E2F transcription factors (16, 17). It possesses two amino acid substitutions, R461E and K542E (R467 and K548 in human numbering), that was designated *Rb1*^*G*^ for simplicity. Surprisingly, these mice had few developmental defects and most animals lived a full, cancer free life span (17). Because they are viable, these mice offer the opportunity to study the physiological scenarios where RB-E2F transcriptional control is most critical. We prepared mouse embryonic fibroblasts from wild type, *Rb1*^*G/G*^, *Rb1*^*−/−*^ animals for comparison of E2F target gene levels and cell cycle control characteristics. Following serum deprivation, relative transcript levels of typical E2F target genes were determined and this demonstrates that *Rb1*^*G/G*^ and *Rb1*^*−/−*^ mutants have similar expression levels, and they are elevated in comparison with wild type controls (Fig. 1A). Following restimulation with serum, BrdU incorporation and flow cytometry were used to determine the proportion of cells undergoing DNA synthesis at each time point (Fig. 1B). This revealed that *Rb1*^*G/G*^ and *Rb1*^*−/−*^ cells also share the same premature cell cycle entry phenotype in response to growth signals. Lastly, we subjected proliferating populations of MEFs to ionizing radiation, followed by BrdU incorporation and detection by flow cytometry. This treatment induced cell cycle arrest and it revealed that *Rb1*^*G/G*^ and *Rb1*^*+/+*^ cells behave similarly and cease proliferation in response to radiation, whereas *Rb1*^*−/−*^ knock out cells are unable to fully respond. These experiments indicate that the *Rb1*^*G*^ mutation compromises some aspects of RB function in cell cycle control in that cells bearing this mutation are defective for E2F transcriptional repression and for controlling cell cycle entry. However, cell cycle exit in response to stimuli such as DNA damage is normal in *Rb1*^*G/G*^ cells suggesting the effects of compromised RB-E2F transcriptional control are dependent on the context of the cell cycle decision. Since we can readily obtain viable *Rb1*^*G/G*^ mice, we crossed them with other strains of mice to challenge *Rb1*^*G*^ cell cycle control defects, or other misregulated functions, and determine their contribution to cancer susceptibility.

**Figure 1:**
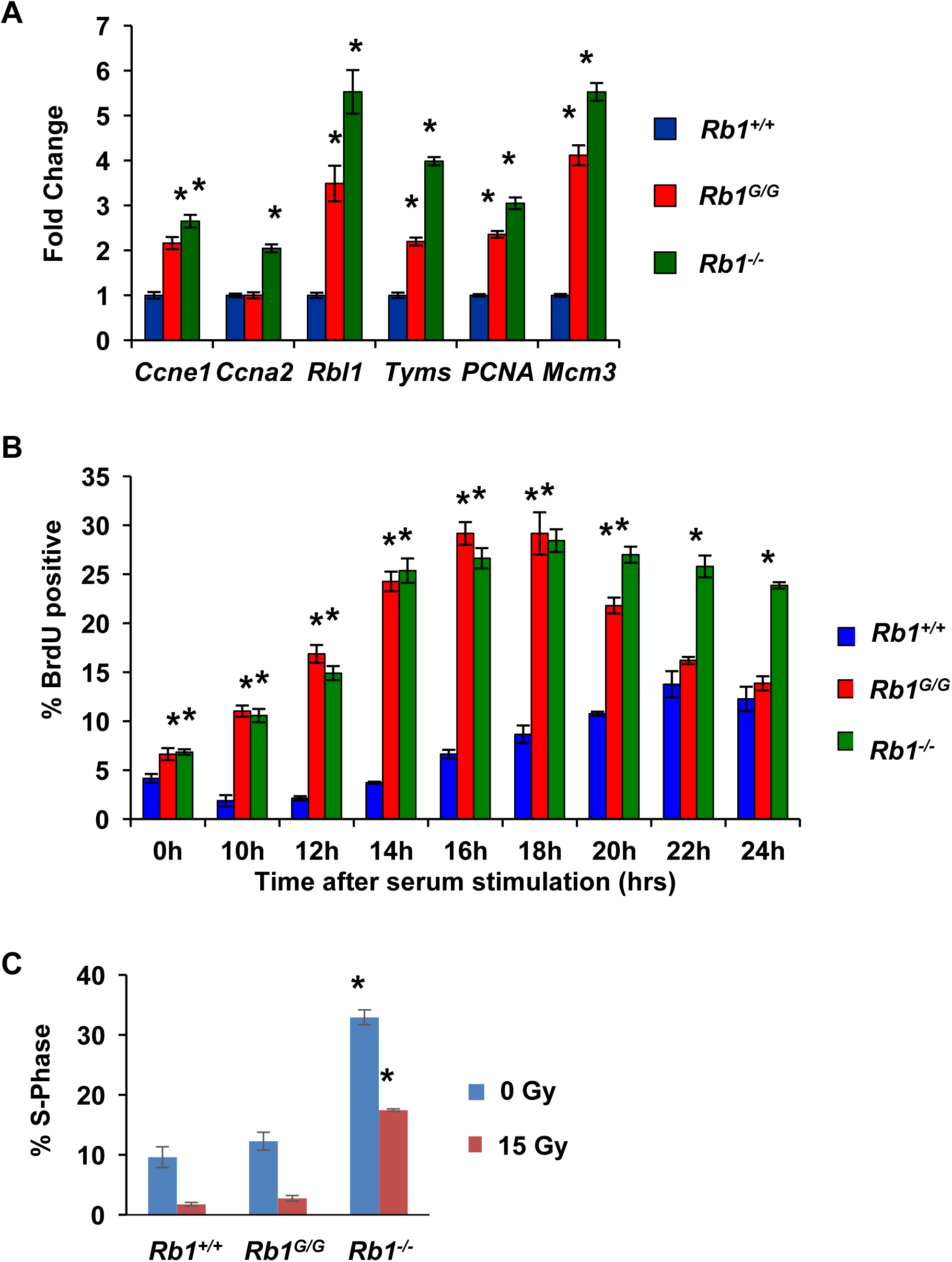
RB-E2F transcriptional control delays cell cycle entry but is dispensable for DNA damage induced arrest. (A) Mouse embryonic fibroblasts (MEFs) were serum starved for 60 hours. Relative transcript levels were determined for known E2F target genes from wild type, *Rb1*^*−/−*^, and *Rb1*^*G/G*^ MEFs. (B) Following serum starvation, MEFs were re-stimulated with serum for the indicated time-course. Cultures were labeled with BrdU for 2 hours before harvesting, staining, and flow cytometry. The percentage of BrdU positive cells is shown for each time point. (C) Proliferating cultures of the indicated genotypes of cells were irradiated with 15 Gy. 48 hours later cells were pulse labeled with BrdU and analyzed by flow cytometry. The percentage of S-phase cells is shown for each. For all experiments, measurements represent the mean of three biological replicates and error bars are one standard deviation. An asterisk indicates a statistical difference from wild type (*t*-test, *p*<0.05).

### The *Rb1*^*G*^ mutation enhances cancer susceptibility of *Trp53*^*−/−*^ mice

Human *RB1* mutations in cancer are frequently accompanied by mutations in *TP53* (25), and cancer phenotypes of *Trp53*^*−/−*^ mice are exacerbated by even a single *Rb1* mutant allele (26, 27). For this reason we crossed *Rb1*^*G/G*^ mice with *Trp53*^*−/−*^ mice. We obtained compound mutant mice within the statistically expected range (Table 1), although we note that *Trp53*^*−/−*^ genotypes appear underrepresented and this may be explained by a previous discovery of partially penetrant neural tube closure defects in embryogenesis (28). Overall, the addition of the *Rb1*^*G*^ mutation does not exacerbate this phenotype, suggesting there are likely no strong developmental phenotypes caused by the combination of these alleles.

**Table 1:** Frequency of generating *Rb1*^*G*^; *Trp53*^*−/−*^ compound mutant mice. The indicated genotypes of mice were crossed and all resulting progeny were genotyped. The number of live animals observed at two weeks of age is indicated for each genotype and the expected number based on Mendelian inheritance is indicated. * denotes significance as determined by chi-squared test.

| | $Rb1^{G/+}; Trp53^{+/-}$ x<br>$Rb1^{G/+}; Trp53^{+/-}$ | |
| --- | --- | --- |
|  | Observed | Expected |
| $Rb1^{+/+}; Trp53^{+/+}$ | 33* | 19 |
| $Rb1^{+/+}; Trp53^{+/-}$ | 44 | 38 |
| $Rb1^{+/+}; Trp53^{-/-}$ | 13 | 19 |
| $Rb1^{G/+}; Trp53^{+/+}$ | 45 | 38 |
| $Rb1^{G/+}; Trp53^{+/-}$ | 80 | 76 |
| $Rb1^{G/+}; Trp53^{-/-}$ | 19* | 38 |
| $Rb1^{G/G}; Trp53^{+/+}$ | 20 | 19 |
| $Rb1^{G/G}; Trp53^{+/-}$ | 39 | 38 |
| $Rb1^{G/G}; Trp53^{-/-}$ | 12 | 19 |
| Total | 305 |  |

We aged cohorts of control *Trp53*^*−/−*^ mutant mice and *Rb1*^*G/G*^; *Trp53*^*−/−*^ animals to determine their susceptibility to cancer. This revealed that *Rb1*^*G/G*^; *Trp53*^*−/−*^ mutant mice experienced a significantly shorter disease free survival compared to *Trp53*^*−/−*^ controls (Fig. 2A). *Trp53*^*−/−*^ knock out mice are known to succumb primarily to thymic lymphomas and sarcomas (26), and we observed both in our control cohort of *Trp53*^*-/ -*^mice. *Rb1*^*G/G*^; *Trp53*^*−/−*^ mice exclusively developed thymic lymphomas and typical thymus morphology and tumor histology are shown (Fig. 2B). This data suggests that combining the *Rb1*^*G*^ mutation with *Trp53* deficiency does not appreciably alter the type of cancer that develops in these mice, but rather accelerates tumor formation. One interpretation of this data is that accelerated cell cycle entry seen in cells from *Rb1*^*G/G*^ mice contributes to cancer susceptibility in *Trp53*^*−/−*^ mutants because it complements a DNA damage induced cell cycle arrest defect, or other cancer relevant deficiency, that is inherent to *Trp53* knock outs. Consistent with this interpretation, we have previously reported that *Rb1*^*G/G*^; *Cdkn1b*^*−/−*^ mice (ie. RB and p27 double mutant) have a synthetic cancer phenotype in which the combined mutation creates deficiency in DNA damage induced cell cycle exit (18).

**Figure 2:**
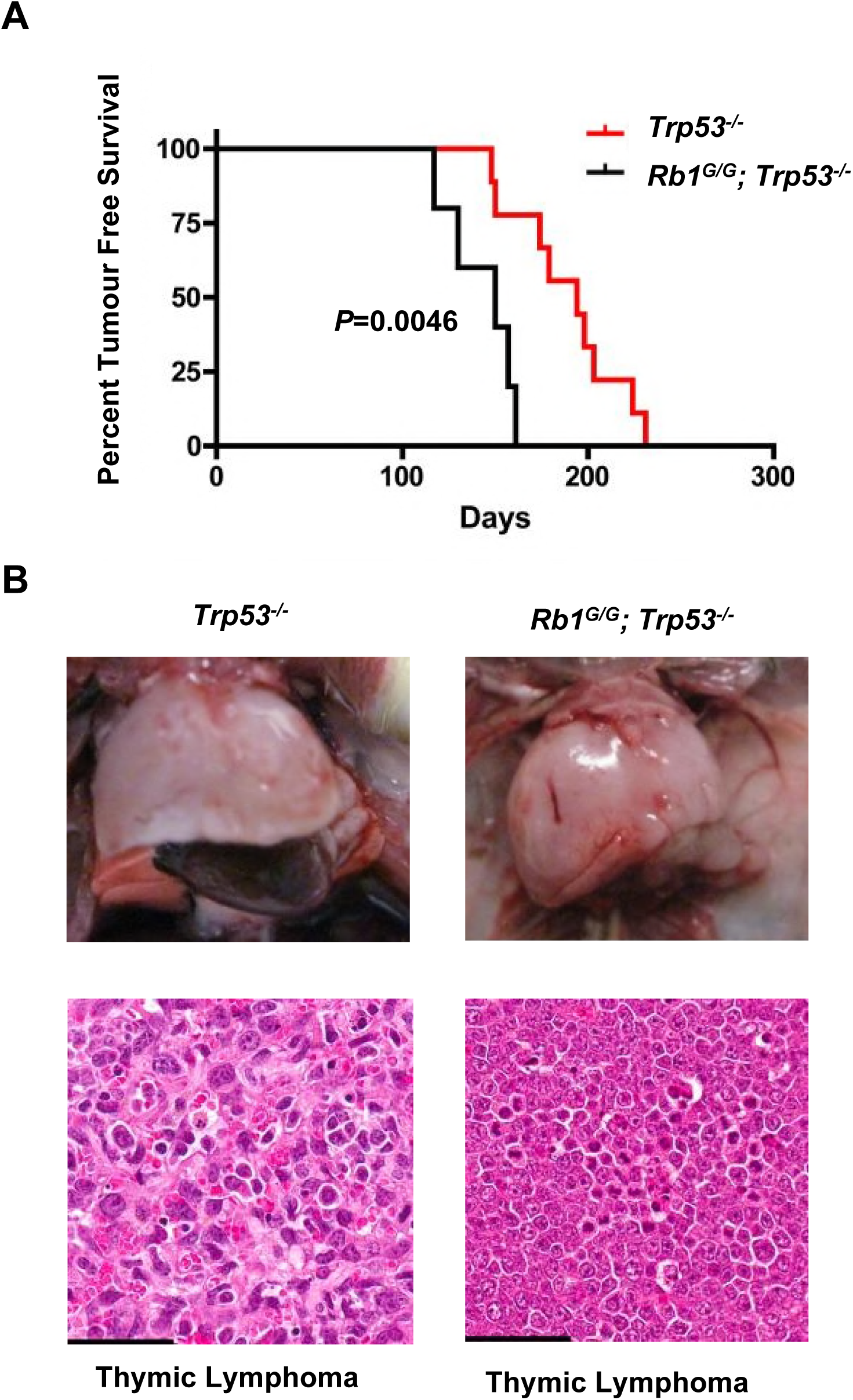
The *Rb1*^*G*^ mutation enhances cancer susceptibility of *Trp53*^*−/−*^ mice. (A) Kaplan-Meier analysis of tumor-free survival of the indicated genotypes. Mice were monitored until their natural endpoint and those having tumors are shown. *Rb1*^*G/G*^; *Trp53*^*−/−*^ (150 days, n=5), *Trp53*^*−/−*^ (194 days, n=9) are statistically significant from one another using the log-rank test (p=0.0046). (B) Whole mount and H&E analysis of thymic lymphomas found in both *Rb1*^*G/G*^; *Trp53*^*−/−*^ and *Trp53*^*−/−*^ mice. Scale bars are equal to 50μm.

### Compound mutant *Rb1*^*G/G*^; *Cdkn1a*^*−/−*^ mice do not succumb to pituitary tumors

Based on the previous observation that *Rb1*^*G/G*^; *Trp53*^*−/−*^ mice have a shorter latency to tumor formation than *Trp53*^*−/−*^ controls, and that *Rb1*^*G/G*^; *Cdkn1b*^*−/−*^ mutations cause similar cell cycle control defects and are also cancer prone, we crossed *Rb1*^*G/G*^ mice with p21 deficient animals. As with the cross to *Trp53*^*−/−*^ mice, *Rb1*^*G/G*^; *Cdkn1a*^*−/−*^ mice were obtained at the expected Mendelian ratios suggesting that the combination of these mutations does not compromise early development of these animals (Table 2).

**Table 2:** Frequency of generating *Rb1*^*G*^; *Cdkn1a*^*−/−*^ compound mutant mice. The indicated genotypes of mice were crossed and all resulting progeny were genotyped. The number of live animals oberved at two weeks of age is indicated for each genotype and the expected number based on Mendelian inheritance is indicated. * denotes significance as determined by chi-squared test.

| | $Rb1^{G/+}; Cdkn1a^{+/-}$ x $Rb1^{G/+}; Cdkn1a^{+/-}$ | |
| --- | --- | --- |
|  | Observed | Expected |
| $Rb1^{+/+}; Cdkn1a^{+/+}$ | 17 | 21 |
| $Rb1^{+/+}; Cdkn1a^{+/-}$ | 64* | 43 |
| $Rb1^{+/+}; Cdkn1a^{-/-}$ | 26 | 21 |
| $Rb1^{G/+}; Cdkn1a^{+/+}$ | 29* | 43 |
| $Rb1^{G/+}; Cdkn1a^{+/-}$ | 82 | 86 |
| $Rb1^{G/+}; Cdkn1a^{-/-}$ | 39 | 43 |
| $Rb1^{G/G}; Cdkn1a^{+/+}$ | 19 | 21 |
| $Rb1^{G/G}; Cdkn1a^{+/-}$ | 37 | 43 |
| $Rb1^{G/G}; Cdkn1a^{-/-}$ | 29 | 21 |
| Total | 342 |  |

We isolated fibroblasts from *Rb1*^*G/G*^; *Cdkn1a*^*−/−*^ mice to test their cell cycle arrest properties in response to ionizing radiation (Fig. 3A). The indicated genotypes of cells were irradiated and BrdU incorporation was used to determine the percentage of cells in S-phase after two days. As expected *Cdkn1a*^*−/−*^ cells were defective for arrest and *Rb1*^*G/G*^; *Cdkn1a*^*−/−*^ compound mutants were similarly unable to arrest in response to irradiation. This indicates that cells in *Rb1*^*G/G*^; *Cdkn1a*^*−/−*^ mice possess mutations that compromise cell cycle entry and exit, akin to *Rb1*^*G/G*^; *Trp53*^*−/−*^ and *Rb1*^*G/G*^; *Cdkn1b*^*−/−*^ mice. To investigate cancer predisposition we generated cohorts of *Cdkn1a*^*−/−*^ and *Rb1*^*G/G*^; *Cdkn1a*^*−/−*^ mutants, as lack of cancer susceptibility of *Cdkn1a* knock out mice has only been reported for young mice (21, 29). Surprisingly, compound mutant mice in both cohorts lived a similar lifespan, and neither has a meaningful cancer predisposition (Fig. 3B-C). This is in clear contrast to *Rb1*^*G/G*^; *Cdkn1b*^*−/−*^ whose cancer free survival data is shown in Figure 3C for comparison purposes. Because *Rb1*^*G/G*^; *Cdkn1b*^*−/−*^ mice develop pituitary tumors with complete penetrance, we also inspected whole mount views of pituitary glands to search for even subtle signs of overgrowth in *Rb1*^*G/G*^; *Cdkn1a*^*−/−*^ mice at survival endpoints (Fig. 3D). We were unable to find pituitary glands in any *Cdkn1a*^*−/−*^ or *Rb1*^*G/G*^; *Cdkn1a*^*−/−*^ mice that were enlarged, and nothing resembling the large pituitary masses observed in *Rb1*^*G/G*^; *Cdkn1b*^*−/−*^ mice (Fig. 3D). This is an important observation as *Rb1*^*+/−*^; *Cdkn1a*^*−/−*^ mice are susceptible to pituitary tumors and have similar survival as for *Rb1*^*G/+*^; *Cdkn1b*^*−/−*^ animals (18, 29), this indicates that the *Rb1*^*G*^ mutation cooperates preferentially with p27 loss.

**Figure 3:**
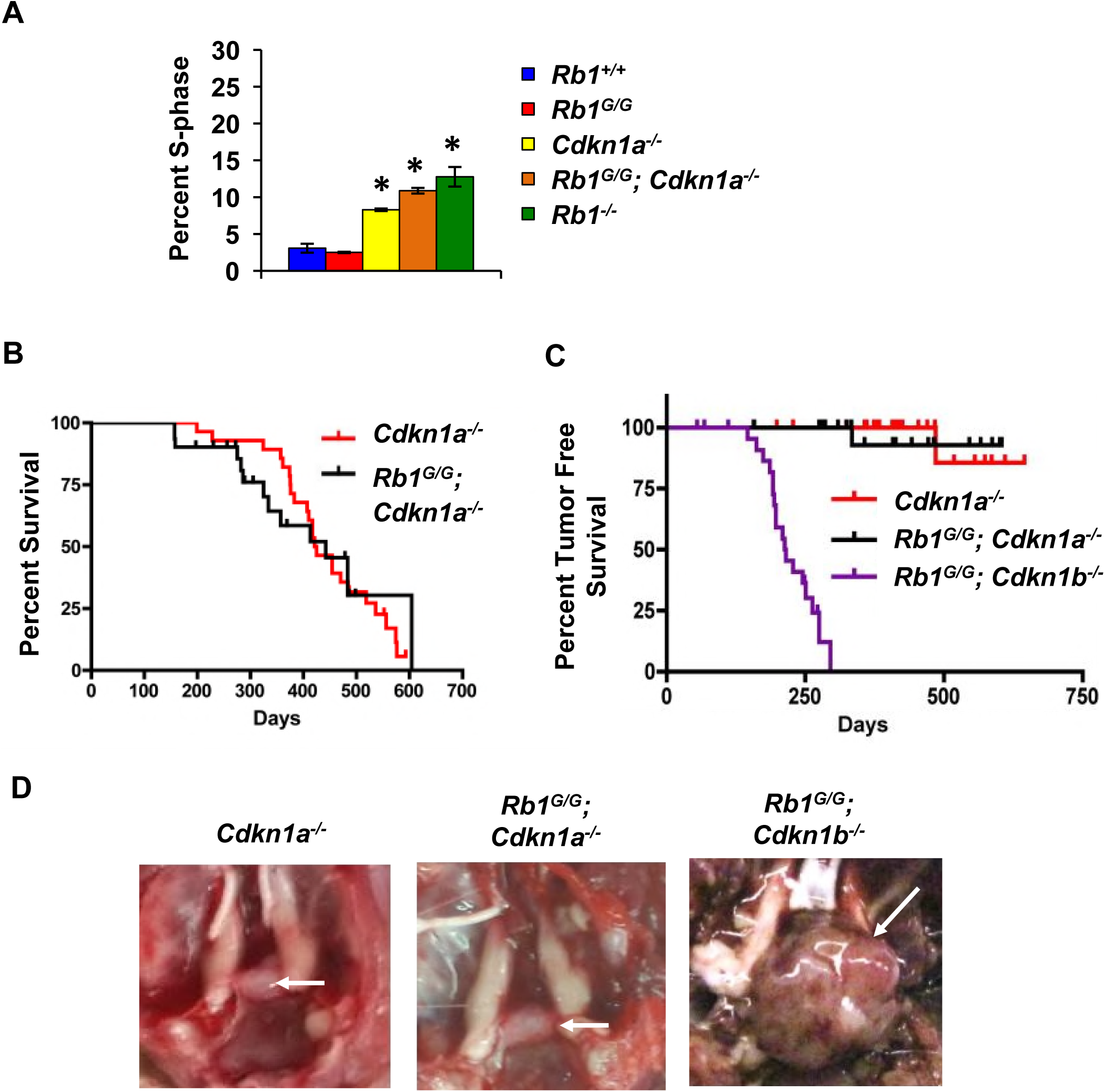
Combining the *Rb1*^*G*^ mutation with *Cdkn1a* deficiency does not create cancer susceptibility. (A) Propidium iodide and BrdU staining of MEFs following 15 Gy of ionizing radiation. S-phase was determined by BrdU incorporation and flow cytometry. An average of 3 replicates are shown, error bars indicate one standard deviation. An asterisk indicates a significant difference from wild type (p<0.05 using a t-test). (B) Kaplan-Meier analysis of overall survival of the indicated genotypes of mice. Mice were monitored until protocol endpoints. The mean survival of *Rb1*^*G/G*^; *Cdkn1a*^*−/−*^ (419 days, n=25) and *Cdkn1a*^*−/−*^ (442 days, n=24) mice are not statically different from one another using the log-rank test (p=0.9059). (C) Kaplan-Meier analysis of tumor-free survival of the indicated genotypes. *Rb1*^*G/G*^; *Cdkn1a*^*−/−*^ and *Cdkn1a*^*−/−*^ are not statically significant from one another using the log-rank test (p=0.7919). Previously determined cancer free survival of *Rb1*^*G/G*^; *Cdkn1b*^*−/−*^ mice is shown for comparison. (D) Whole mount images of pituitaries of aged *Cdkn1a*^*−/−*^ and *Rb1*^*G/G*^; *Cdkn1a*^*−/−*^ animals demonstrate normal anatomical structure. A pituitary tumor from an *Rb1*^*G/G*^; *Cdkn1b*^*−/−*^ mouse is shown for comparison. Pituitary glands/tumors are indicated by white arrows.

Deficiency for *Trp53* and CDK inhibitor proteins undeniably alters cell cycle control. Mutant mice that combine these mutations with the defects present in *Rb1*^*G/G*^ mice exacerbate cell cycle control defects further through accelerated G1-S transition. However, the lack of cancer predisposition in *Rb1*^*G/G*^; *Cdkn1a*^*−/−*^ mice suggests that these deficiencies aren’t cancer enabling on their own. Possible explanations for why *Trp53* and *Cdkn1b* mutations accelerate or enable cancer in combination with the *Rb1*^*G*^ mutation when *Cdkn1a* knock out does not will be discussed below.

### The *Rb1*^*G*^ mutation does not enhance *Kras* driven proliferation

One shortcoming of investigating cancer predisposition in mutant mice that uniformly possess loss of function mutations is that a critical initiating event may be missing. For this reason we also crossed *Rb1*^*G/G*^ mutants with mice expressing an oncogenic form of Kras in order to stimulate proliferation and provide aberrant survival signals. To accomplish this we utilized the latent LSL-KrasG12D mutant that has previously been reported (23). In order to be consistent with previous experiments we utilized the UBC9-Cre-ERT2 transgene and mild tamoxifen delivery so that the *Kras*^*G12D*^ allele is expressed in some cells in potentially many tissues.

We generated cohorts of *LSL-Kras*^*G12D*^; *UBC-Cre-ERT2* and *Rb1*^*G/G*^; *LSL-Kras*^*G12D*^; *UBC-Cre-ERT2* mice. Mice were given two doses of tamoxifen to generate sporadic activated Kras. In all cases, these mice developed hyperplasia originating in the upper gastrointestinal tract. Histologically we have classified these as squamous papillomas and they often extend into the mouths and stomachs of affected mice (Fig. 4A). This phenotype was not altered by the *Rb1*^*G*^ mutation (Fig. 4B), nor was the overall survival of these mice different between cohorts (Fig. 4C). This experiment suggests that the *Rb1*^*G*^ allele doesn’t accelerate the growth of squamous papillomas in compound mutant mice, nor does it cooperate with activated Kras to transform these lesions into carcinomas. However, there are some considerations that should be taken into account when interpreting this data and they are discussed below.

**Figure 4:**
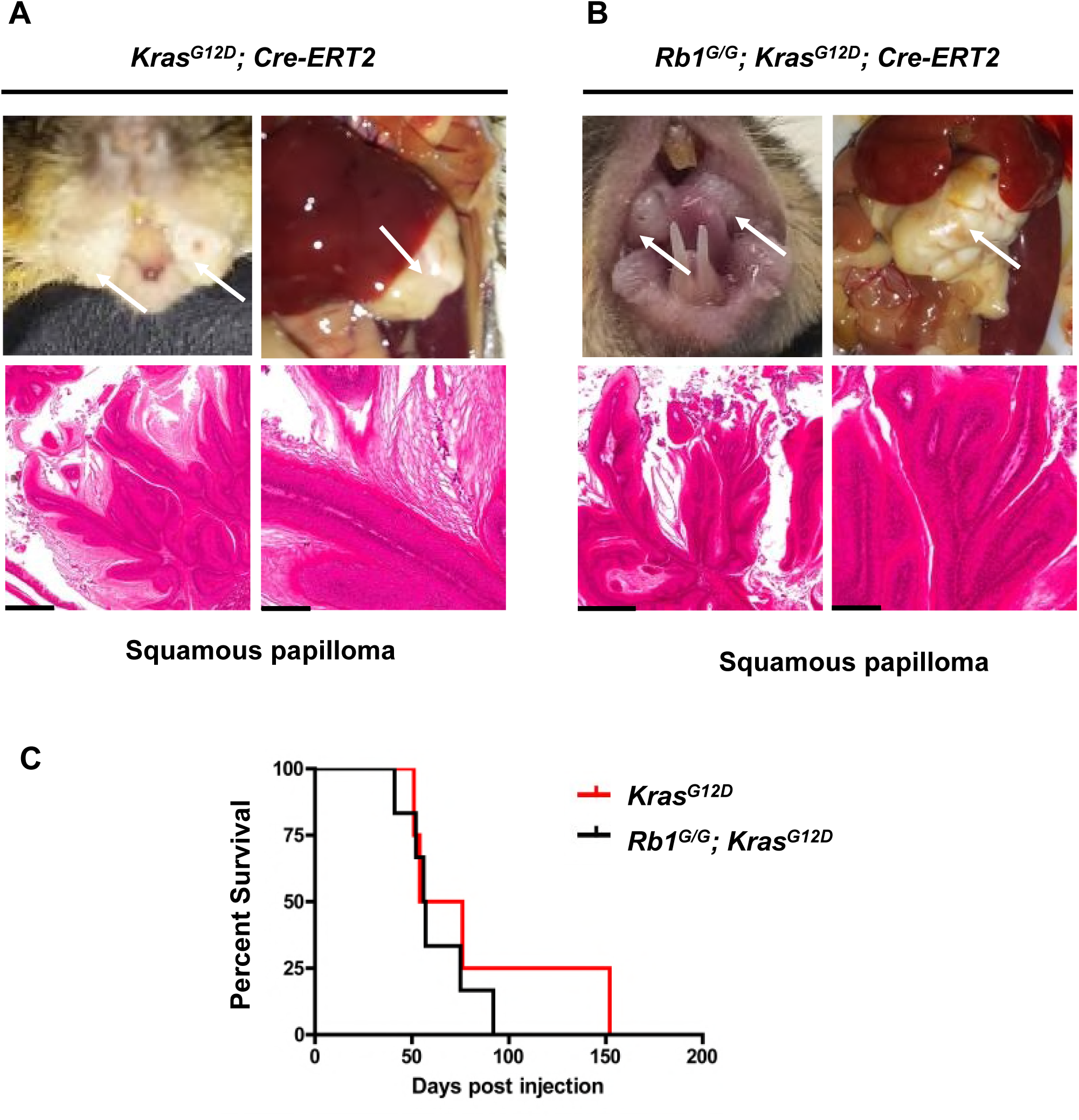
The *Rb1*^*G*^ mutation does not modify cancer susceptibility of Kras^G12D^ expressing mice. (A) Gross morphology of tissue growths that are evident in the mouths and stomachs of tamoxifen treated *UBC-Cre-ERT2; Kras*^*G12D*^ mice (top left and right respectively). (B) Morphology of growths in *UBC-Cre-ERT2; Rb1*^*G/G*^; *Kras*^*G12D*^ mice (top left and right). Hematoxylin and Eosin staining (lower panels of A and B) show squamous papillomas found in the mouths and stomachs of control *Kras*^*G12D*^ and *Rb1*^*G/G*^; *Kras*^*G12D*^ animals. Scale bars represent 400μm for low power histology (left) and 100μm for high power (right). (C) Kaplan-Meier analysis of survival of the indicated genotypes. Mice were monitored until natural endpoint. *Rb1*^*G/G*^; *Kras*^*G12D*^ (56.5 days post injection) (n=6) and *Kras*^*G12D*^ mice (65 days post injection) (n=4) are not statistically different from one another using the log rank test (p=0.475).

## Discussion

In this report, we aimed to investigate the role of RB-E2F transcriptional control in isolation from other RB family E2F regulation using cancer susceptibility of *Rb1*^*G/G*^ mice as a simple read out. We have shown that the *Rb1*^*G*^ mutation exacerbates cancer development in *Trp53*^*−/−*^ mice, but has essentially no effect on tumor-free survival in the presence of oncogenic Kras^G12D^, or from loss of p21 (*Cdkn1a*^*−/−*^). In conjunction with prior results that *Rb1*^*G/G*^; *Cdkn1b*^*−/−*^ animals succumb to pituitary tumors (18), and that *Rb1*^*+/−*^; *Cdkn1a*^*−/−*^ mice have accelerated pituitary tumorigenesis (29), these genetic crosses provide two simple conclusions relating to RB function in cell cycle and cancer susceptibility. The first is that compromised RB-E2F transcriptional regulation is not uniformly cancer promoting, even though it potently contributes in some crosses. Cancer susceptibility in these crosses does not correlate with obvious cell cycle control defects. The second conclusion is that loss of p27 contributes to RB-E2F driven cancer because of functions it doesn’t share with p21, and these are likely beyond cell cycle control. These conclusions will be discussed further, along with limitations of our study.

Oncogenic Kras^G12D^ causes a signalling cascade that results in both the expression of E2Fs, an increase in Cyclin/CDK complexes, and the suppression of p27 activity (30, 31). From this perspective we expected Kras activation may be more potent when combined with the *Rb1*^*G*^ mutation since *Rb1*^*G*^ cooperates with p27 loss and is defective for regulating E2F transcription factors. However, there are two caveats that temper our interpretation of an inability of the *Rb1*^*G*^ mutation to enhance Kras driven hyperplasia. First, the extremely fast rate at which expression of Kras^G12D^ induces squamous papillomas (median survival was 50 days), suggests that the *Rb1*^*G*^ mutation may not be able to detectably exacerbate these effects in such a short time course. In addition, since these mice uniformly succumb to lesions in the upper gastrointestinal tract, their anatomical location almost immediately triggers protocol endpoints preventing the study of long term effects. Slower developing tumors in other tissues may be more amenable to studying *Rb1*^*G*^ effects. We expect this would require Cre drivers that afford a tissue by tissue approach as a number of studies have used other ubiquitous Cre expression approaches that have resulted in different cancer types (32, 33), or different benign lesions (32), than what we observe with this UBC9 driven CreERT2. For these reasons it is difficult to draw clear mechanistic conclusions from this cancer cross. The other crosses described in this report, and those referenced as comparisons, use constitutive mutant alleles and longer time courses allowing for more definitive conclusions.

Based on the cancer susceptibility data presented in this study, the cancer initiation contexts that benefit from defective RB-E2F regulation are shared by p53 and p27 deficiency, but not p21 deficiency and this leads to the comparison of p27 and p21 functions. Both are members of the CIP/KIP family of CDK inhibitors that influence cell cycle control through the inhibition of cyclin/CDK phosphorylation (34). However, p27 is reported to possess a number of functions related to lineage commitment and cell migration control that are distinct from p21 (35). For example, p27 is able to regulate the expression of the stem cell transcription factor *Sox2* through the regulation of the SRR2 enhancer (36). Similarly, RB also occupies *Sox2* at the SRR2 enhancer, and the *Rb1*^*G*^ allele is defective for transcriptional repression of *Sox2* and fails to be recruited to this location (37). This example of a context where RB-E2F transcriptional control may play a tumor suppressive role is particularly relevant because pituitary tumorigenesis in *Rb1*^*+/−*^mice is dependent on *Sox2* (37). In addition, p53 deficiency also contributes to *SOX2* misexpression and lineage plasticity in prostate cancer and this effect is most pronounced when combined with *RB1* mutations (38). Thus, scenarios such as this one may explain molecular connections between RB-E2F regulation and contexts where its loss contributes to cancer. Surprisingly, cell cycle control defects alone are unlikely to explain the contexts where CDK inhibitors and RB-E2F transcriptional regulation are most critical, as this example illustrates.

This study offers a fresh look at RB’s role as a tumor suppressor as our work shows that defects in its best known function, regulation of E2F transcription, is not uniformly cancer promoting. We expect that with new CDK4/6 inhibitors that require RB for efficacy arriving in the clinic, understanding the contexts of RB function will be important to ensure that these inhibitors are given to patients with the best chance of response.

## Acknowledgements

The authors wish to thank colleagues in the London Regional Cancer Program for frequent discussions and encouragement during the course of this study. MJT was a members of the CIHR Strategic Training Program in Cancer Research (CaRTT), and a recipient of funding from OGS. FAD is the Wolfe Senior Fellow in Tumor Suppressor Genes at Western University.

